# Fluorescence lifetime imaging microscopy for metabolic analysis of LDHB inhibition in triple negative breast cancer

**DOI:** 10.1101/2025.01.13.632864

**Authors:** A. Galloway, B. Ter Hofstede, Alex J. Walsh

## Abstract

Triple-negative breast cancer (TNBC) is an aggressive subtype of breast cancer with no targeted treatments currently available. TNBC cells participate in metabolic symbiosis, a process that optimizes tumor growth by balancing metabolic processes between glycolysis and oxidative phosphorylation through increased activity by the enzyme lactate dehydrogenase B (LDHB). Metabolic symbiosis allows oxidative cancer cells to function at a similar rate as glycolytic cancer cells, increasing overall metabolic activity and proliferation. Here, fluorescence lifetime imaging microscopy (FLIM) is used to analyze the metabolism of TNBC cells with inhibition of LDHB using a multiphoton microscope to measure the fluorescent lifetimes of two metabolic coenzymes, NAD(P)H and FAD. LDHB is inhibited via an indole derivative known as AXKO-0046 in varying concentrations. Understanding how TNBC cell metabolism changes due to LDHB inhibition will provide further insight into metabolic symbiosis and potential new TNBC treatment options.

## 1. Introduction

Triple negative breast cancer (TNBC) is an aggressive subtype of breast cancer that lacks estrogen receptors, progesterone receptors, and human epidermal growth factor receptor 2 (HER2) amplification within their tumor cells [1]. TNBC typically has a poor prognosis due to the invasiveness of the cancer and limited treatment options. To better understand how TNBC works and find effective treatments, TNBC cellular metabolism needs to be better understood.

Cellular metabolism is the process of consuming and creating energy in the form of adenosine triphosphate (ATP) by complicated processes such as glycolysis and oxidative phosphorylation (OXPHOS) [2]. Glycolysis yields ATP as a result of converting glucose to pyruvate [3]. In aerobic conditions, pyruvate produced from glycolysis is used in the tricarboxylic acid (TCA) reactions which then leads to OXPHOS and yields ATP but in a higher amount than seen in glycolysis alone [4]. While most healthy cells prefer OXPHOS when oxygen is available, some cancer cells prefer glycolysis despite oxygen abundance, known as the Warburg effect [5]. In most TNBC tumors, the cellular population is made up of both glycolytic and oxidative cancer cells, meaning these metabolic pathways coexist, leading to rapid cell growth characteristic of tumors [6]. Because both pathways require glucose but typically the tumor microenvironments do not contain an abundance of glucose, TNBC cells utilize their metabolic plasticity and rewire their metabolic reactions to optimize cellular proliferation [7]. This process is known as metabolic symbiosis and is dependent on lactate [8]. Instead of OXPHOS depending on glycolysis to convert glucose to pyruvate for the TCA reactions, lactate as a byproduct of glycolysis is converted back to pyruvate via lactate dehydrogenase B (LDHB), forgoing the need for oxidative cells to initially depend on glucose [8]. The reaction controlled by LDHB and its significant upregulation in cancer is what makes metabolic symbiosis possible [9].

Interfering with TNBC’s metabolic symbiosis may be a promising therapeutic approach to slow cancer growth. If the collaboration between oxidative and glycolytic cancer cells is interrupted, theoretically the rapid cell growth of tumors will decrease due to a less efficient use of glucose. While the role of LDHA in cancer metabolism is known, the function and contribution of LDHB to cancer growth is not well understood [10]. This research aims to characterize how cellular metabolism changes following LDHB inhibition. Using a TNBC cell line, MDA-MB-231, LDHB was inhibited chemically using the first known highly selective human LDHB inhibitor, AXKO-0046 [11]. The metabolic changes were observed using fluorescence lifetime imaging microscopy (FLIM), a label-free optical imaging tool used to characterize cellular metabolism by using two fluorescent metabolic coenzymes, nicotinamide adenine dinucleotide (NAD(P)H) and flavin adenine dinucleotide (FAD) [12]. The optical redox ratio, defined here as the NAD(P)H intensity divided by the summed intensity of NAD(P)H and FAD, was used to measure changes in cellular metabolism, with a higher redox ratio corresponding to increased glycolysis or reduced OXPHOS and a lower redox ratio corresponding to increased OXPHOS or reduced glycolysis [12]. Label-free imaging technology, such as FLIM, allows for an understanding of how LDHB inhibition changes tumor metabolism without invasive techniques, aiding in the development of TNBC treatments with a focus on metabolic symbiosis.

## 2. Methods

### 2.1 Cell Culture

MDA-MB-231 cells (ATCC #HTB-26) were cultured according to Baek et al. in RPMI-1640 (Gibco) with 5 mM glucose, 10% FBS, and 1x antibiotic-antimycotic solution [14]. Cells were incubated at 37°C, 5% CO_2_, and 95% relative humidity. 48 hours before imaging, MDA-MB-231 cells were plated at a density of approximately 100,000 cells on 35 mm glass-bottom imaging dishes (MatTek). A 1mM stock solution of AXKO-0046 (MedChemExpress #HY-147216) dissolved in DMSO was added to dishes and incubated in varying concentrations (0 μM (control), 0.1 μM, and 1.0 μM) at 1 hour, 12 hour, and 24 hour time points before image acquisition.

### 2.2 Image acquisition

Images were acquired using a multiphoton laser-scanning microscope (Marianas, 3i) with a 100X objective (NA = 1.46, Zeiss). To capture metabolic information, emission filters of 447/60 nm and 550/88 nm were used for NAD(P)H and FAD, respectively. NAD(P)H was excited at 750 nm and FAD was excited at 890 nm. Fluorescence lifetime images were obtained using photomultiplier tube (PMT) detectors and time-correlated single-photon counting (TCSPC) electronics (SPC-150N, Becker & Hickl). About 6 images were captured per dish with a dwell time of 50 μs and an image size of 256×256 pixels. A stage top incubator maintained the cells at 37°C, 5% CO_2_, and 85% relative humidity during imaging sessions.

### 2.3 FLIM data analysis

Fluorescence lifetime information of NAD(P)H and FAD were analyzed by SPCImage (Becker & Hickl). The lifetime value of each cell was calculated based on the decay curve of each pixel which was deconvoluted from the measured IRF of urea crystals, then fitted to a bi-exponential decay model (***Equation 1***) and binned by 9 surrounding pixels (bin=1). The fluorescence intensity as a function of time *t* is represented by *I(t)*. The fractions of the short and long lifetimes are represented by α_1_ and α_2_ respectively while τ_1 &_ τ_2_ represent the average short and long lifetimes, respectively. C accounts for background light. When NAD(P)H is in its free conformation, it has a short lifetime (τ_1_); when bound to a protein, it has a long lifetime (τ_2_). FAD follows the opposite trend where τ_1_ and τ_2_ correspond to the average lifetime of bound FAD and free FAD respectively. To acquire cell-based fluorescence lifetime endpoints, NAD(P)H images were first segmented using CellPose [15]. Next, MATLAB was used to calculate the average optical redox ratio (***Equation 2***) and lifetime features of each segmented cell. Statistical analysis was performed in R. Statistical significance was determined using multiple t-tests between each experimental group and time points.

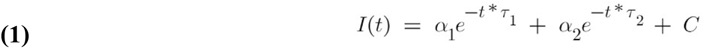

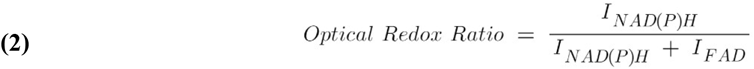

### 2.4 ImageJ Image Analysis

NAD(P)H intensity images were uploaded to ImageJ and thresholded to include contents of the entire cell to then set all background pixels to nan. The corresponding FAD images were uploaded and the optical redox ratio image was created by using the image calculator function. First, the NAD(P)H and FAD images were added and then the NAD(P)H image was divided by the resulting summed image. Finally, the fire LUT was added onto the optical redox ratio images and the images were scaled to include optical redox ratio values of 0.25-1 to show spatial distribution of optical redox ratio.

## 3. Results

Qualitative analysis using ImageJ of the optical redox ratio of MDA-MB-231 cells treated with AXKO-0046 revealed changes in cellular morphology as shown in representative images (Fig. 1). The 1.0 μM groups across all time trials exhibited a circular cell shape compared to the control and 0.1 μM indicating detachment of cells from the culture flask and early stages of apoptosis (Fig. 1) [16]. Additionally, the images are false-colored based on the optical redox ratio value at each pixel, with lower values indicating where a transition from glycolysis to OXPHOS is occurring spatially in the cell (Fig. 1).

**Figure 1:**
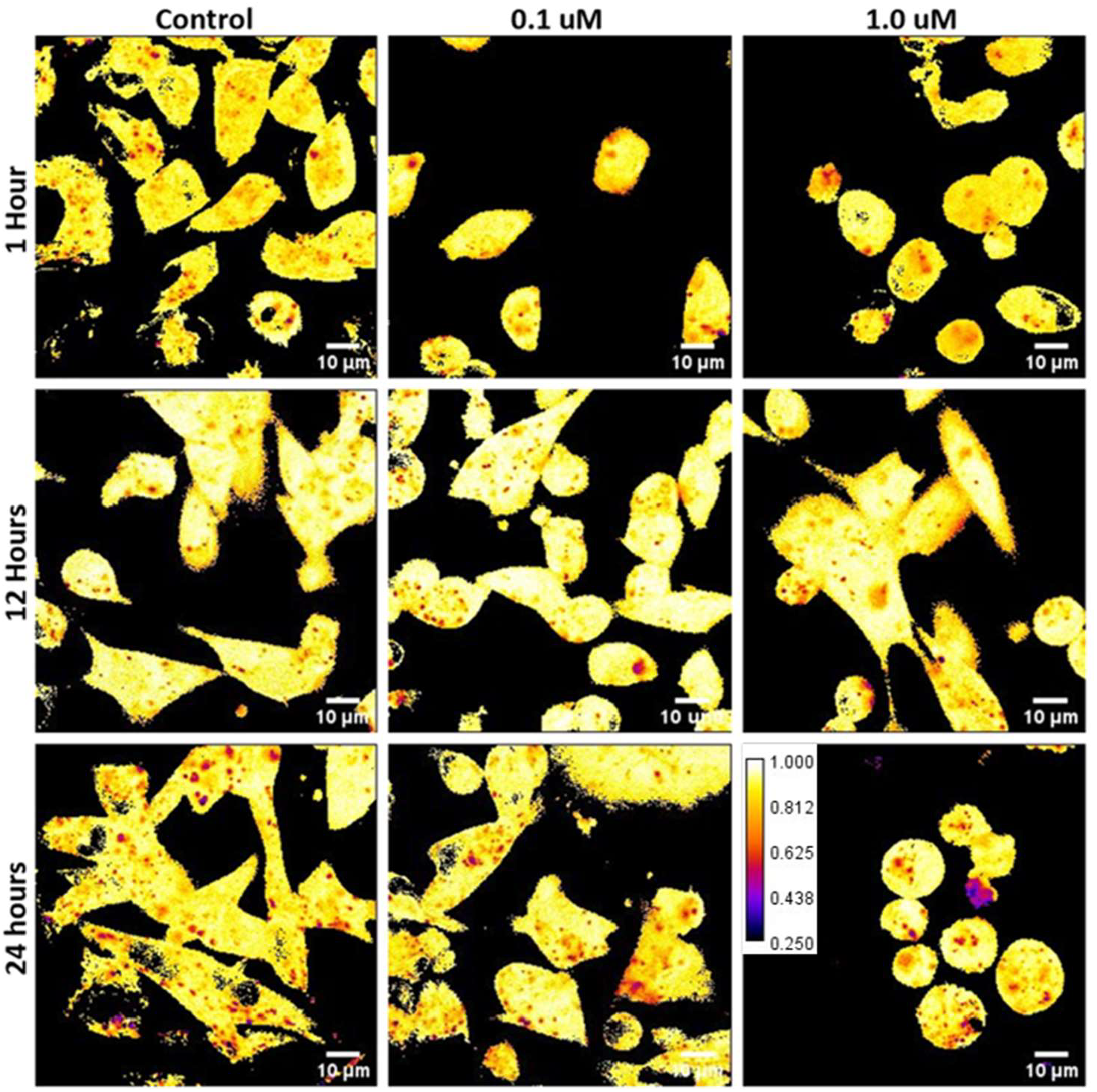
Representative images of optical redox ratio of MDA-MB-231 cells treated with LDHB inhibitor (control, 0.1 μM, 1.0 μM) at 1 hour, 12 hours, and 24 hours after treatment. Optical redox ratio is defined by NAD(P)H/(NAD(P)H + FAD).

Quantitative analysis of the optical redox ratio of MDA-MB-231 cells treated with AXKO-0046 revealed no statistically significant changes as shown in Figure 2. The optical redox ratio was consistent across control and cells treated with 0.1 μM and 1.0 μM AXKO-0046 and across time points of 1, 12, and 24 hours of incubation (Fig. 2).

**Figure 2:**
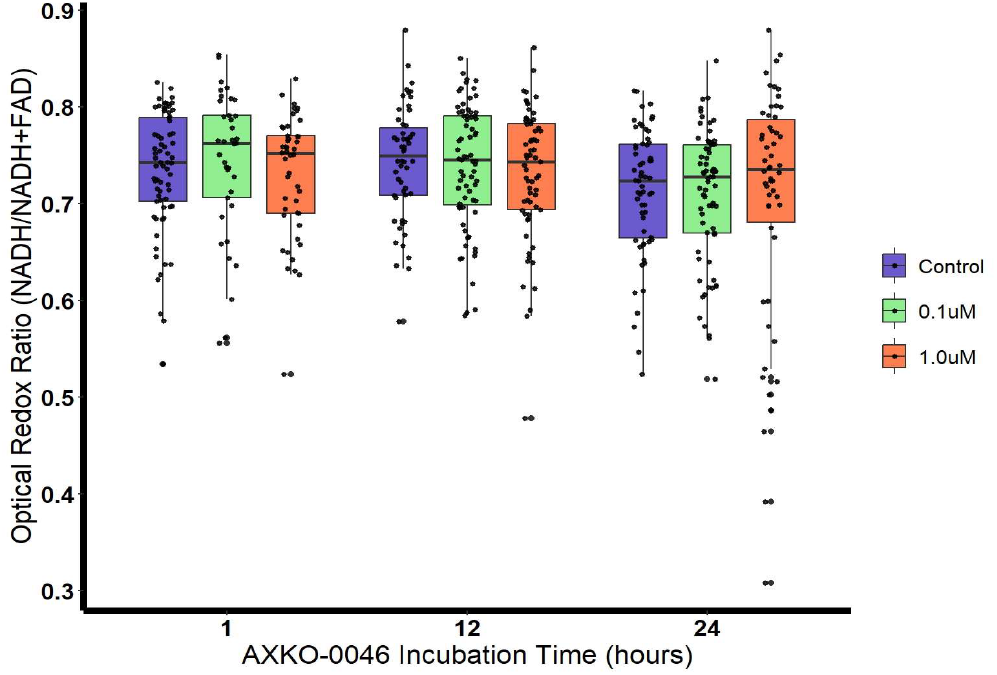
Optical redox ratio of MDA-MB-231 cells after LDHB inhibitor (AXKO-0046) treatment at different concentrations and incubation durations.

Analysis of various FLIM parameters revealed many statistically significant changes upon AXKO-0046 treatment at the different timepoints (Fig. 3). The percentage of free NAD(P)H significantly increased for the 1.0 μM treatment group at 24 hours compared to both the 1 hour and 12 hour 1.0 μM treatment groups (Fig. 3A). Similarly, there was a statistically significant increase of the bound FAD between the 1.0 μM treatment group at 1 hour and 12 hours as well as between the 1.0 μM treatment group at 1 hour and 24 hours (Fig. 3D). At 24 hours, there was a statistically significant difference between both the control and 0.1 μM treatment group and the 0.1 μM and 1.0 μM treatment groups, but no significant changes between the control and 1.0 μM group (Fig. 3D). The average lifetime of free NAD(P)H of the 1.0 μM treatment group at 24 hours significantly increased compared to the control at 24 hours, 0.1 μM treatment group at 24 hours, and the 1.0 μM treatment groups at both 1 and 12 hours (Fig. 3B). The average fluorescent lifetime of bound NAD(P)H follows similar trends to the average fluorescent lifetime of free NAD(P)H (Fig. 3C). For the average fluorescent lifetime of bound FAD, there was a statistically significant increase between the control and 1.0 μM treatment group at 1 hour (Fig. 3E). However, there was a statistically significant decrease in the average fluorescent lifetime of bound FAD in the 1.0 μM treatment groups at 12 and 24 hours when compared to the 1 hour incubation time (Fig. 3E). The average fluorescent lifetime of free FAD followed a similar trend to bound FAD. There was a statistically significant increase in the average fluorescent lifetime between the control and 0.1 μM treatment group as well as the control and 1.0 μM treatment group at the 1 hour time trial (Fig. 3F). However, there was a statistically significant decrease in the average fluorescent lifetime of free FAD in the 1.0 μM treatment groups at 12 and 24 hours when compared to the 1 hour incubation time (Fig. 3F). Lastly, there was also a statistically significant decrease between the 0.1 μM and 1.0 μM treatment groups at the 24 hour time point (Fig. 3F).

**Figure 3:**
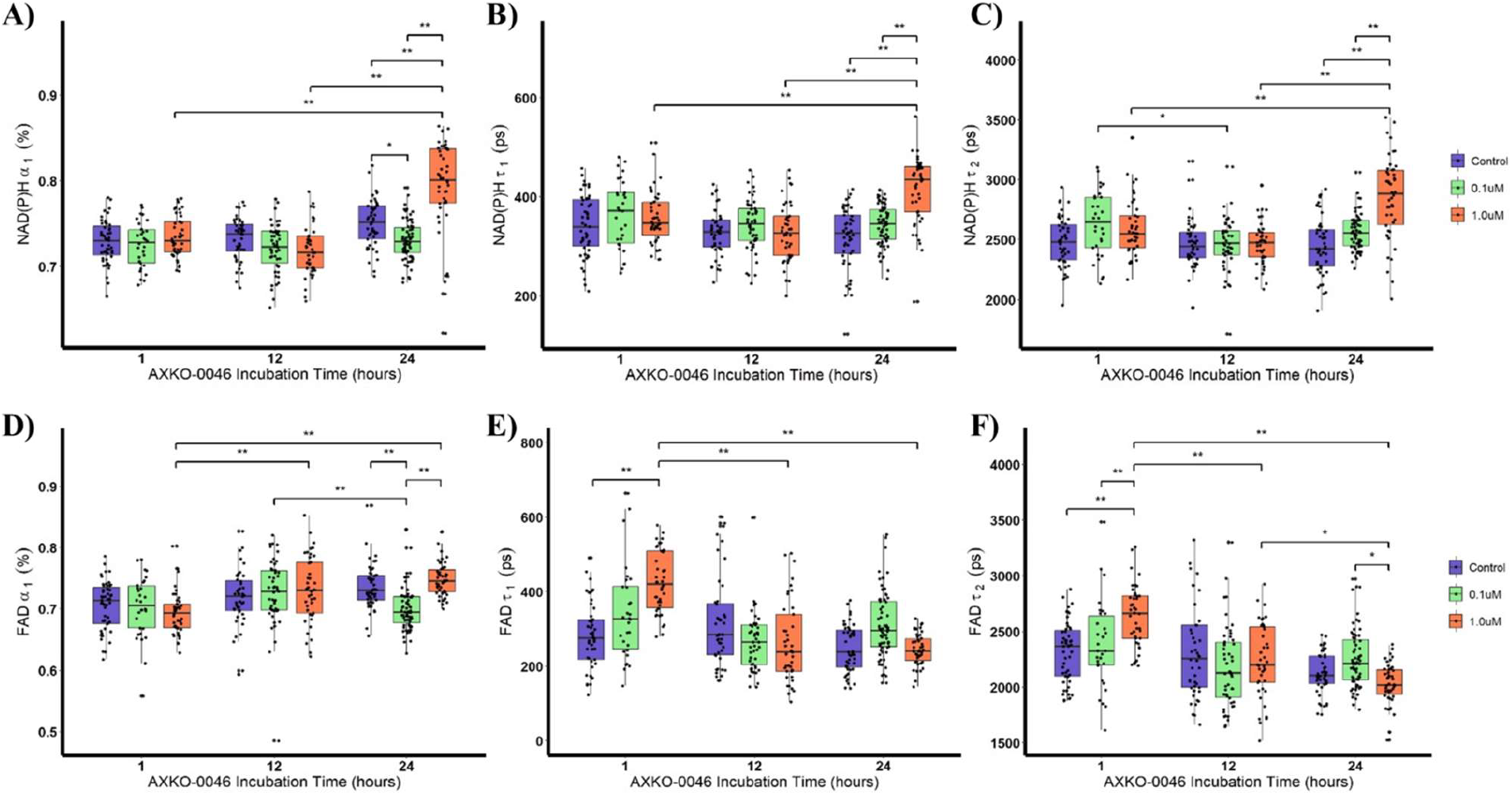
Average NAD(P)H (A-C) and FAD (D-F) fluorescent lifetime endpoints of MDA-MB-231 cells at times (1 hour, 12 hours, and 24 hours) after varying concentrations of AXKO-0046 treatment.

## 4. Discussion

LDH contributes to the progression of cancer, thus better understanding its role could lead to possible cancer treatments. Because triple negative breast cancer overexpresses LDHB and depends on the reaction it catalyzes for cellular metabolism and proliferation, LDHB inhibition may slow cancer progression. To analyze cellular metabolism, FLIM was performed to evaluate the effect of the LDHB inhibitor, AXKO-0046, on the autofluorescence of MDA-MB-231 cells. Because LDHB is inhibited, lactate begins to accumulate due to its inability to convert to pyruvate. When lactate builds up in the cytosol, this causes signaling within the cell to switch from glycolysis to OXPHOS [17]. Although there are no statistically significant changes in the optical redox ratio, shown in *Figure 2*, the 24 hour 1.0 μM treatment group has more variability in its redox ratio, potentially indicating drug-induced changes in metabolism. Further, FLIM analysis revealed a large increase of free NAD(P)H and an increase of both the short and long fluorescent lifetimes of NAD(P)H (Figs. 3A-C), suggesting inhibition of LDHB is changing the metabolism of TNBC cells. Additionally, an increase in both the short and long lifetimes of NAD(P)H has been associated with an increase in OXPHOS, suggesting that AXKO-0046 has interrupted metabolic symbiosis [13]. While the changes in NAD(P)H seem to depend on AXKO-0046 concentration, the changes in FAD appear to be time dependent. Between the 1 hour, 12 hour, and 24 hour time points, the fluorescent lifetimes of FAD follow different trends. This could be due to two things. First, FAD is produced further downstream in OXPHOS than when NAD+ is reduced to NAD(P)H, meaning the effects of LDHB inhibition on FAD could be occurring after the effects on NAD(P)H. Second, the short and long fluorescent lifetimes of FAD is also pH dependent and because of the introduction of a new chemical into the cellular environment, the pH is changing and therefore affecting FAD fluorescent lifetimes.

## 5. Conclusion & Future directions

Because triple negative breast cancer lacks hormones and receptors needed for targeted therapies, chemotherapeutics with a metabolic mechanism of action are favored. Due to the overexpression of LDHB in TNBC and its important role in metabolic symbiosis, inhibition of LDHB serves as a potential treatment option. LDHB inhibition causes more metabolic variability in TNBC, suggesting that it is capable of interrupting metabolic symbiosis. Additionally, LDHB inhibition leads to an increase in both the short and long lifetimes of NAD(P)H, indicating a shift towards OXPHOS. The 24 hour imaging experiment will be repeated on MCF10A cells to examine whether LDHB inhibition negatively affects healthy breast cells. Additionally, viability experiments will be completed on both MDA-MB-231 cells and MCF10A cells to examine how LDHB inhibition affects cellular viability and changes in cellular populations.

## Acknowledgements

This project is supported by funding source NIH NIGMS R35 GM142990 and has been made possible in part by grant number 2022 – 251380 from the Chan Zuckerberg Initiative DAF, an advised fund of Silicon Valley Community Foundation. Additionally, the authors would like to thank Dr. Linghao Hu for assistance with image analysis.

